# Developmental trajectory of cortical somatostatin interneuron function

**DOI:** 10.1101/2024.03.05.583539

**Authors:** Alex Wang, Katie A. Ferguson, Jyoti Gupta, Michael J. Higley, Jessica A. Cardin

## Abstract

GABAergic inhibition is critical to the proper development of neocortical circuits. However, GABAergic interneurons are highly diverse and the developmental roles of distinct inhibitory subpopulations remain largely unclear. Dendrite-targeting, somatostatin-expressing interneurons (SST-INs) in the mature cortex regulate synaptic integration and plasticity in excitatory pyramidal neurons (PNs) and exhibit unique feature selectivity. Relatively little is known about early postnatal SST-IN activity or impact on surrounding local circuits. We examined juvenile SST-INs and PNs in mouse primary visual cortex. PNs exhibited stable visual responses and feature selectivity from eye opening onwards. In contrast, SST-INs developed visual responses and feature selectivity during the third postnatal week in parallel with a rapid increase in excitatory synaptic innervation. SST-INs largely exerted a multiplicative effect on nearby PN visual responses at all ages, but this impact increased over time. Our results identify a developmental window for the emergence of an inhibitory circuit mechanism for normalization.

## Introduction

Neural circuits in the cerebral cortex are comprised of distinct populations of excitatory glutamatergic pyramidal neurons (PNs) and diverse inhibitory GABAergic interneurons (INs). INs are key regulators of adult cortical circuit function, restricting the timing and amplitude of excitatory output from PNs. In addition to their roles in mature circuits, INs are also critical to the proper development of mature cortical circuits. However, the precise roles of distinct subpopulations of INs in regulating cortical circuit development remain largely unclear.

Recent work has highlighted the unique roles of somatostatin-expressing (SST), dendrite-targeting INs in cortical circuit function. SST-INs regulate synaptic integration in PNs ^1–4^ and influence calcium-dependent dendritic plasticity ^5–7^. In primary visual cortex (V1), SST-INs exhibit strong visual responses that are broadly tuned for stimulus size, orientation, and spatial frequency ^8–12^. SST-INs in layer 2/3 receive extensive horizontal excitatory inputs, but little feedforward input ^1,13^, and make promiscuous synapses on local PNs ^14,15^. These INs are unique in exhibiting strong responses to large visual stimuli ^8,16^, potentially leading to surround suppression of local PNs ^13^ and long-range coordination of visually evoked activity ^17^. Moreover, SST-IN activity is robustly modulated by changes in behavioral state, such as arousal and locomotion ^8,16,18^. SST-IN activity increases during locomotion^11^, leading to a locomotion-induced increase in SST-IN visual response gain ^8,16,19^ that may contribute to state-dependent visual perceptual performance via inhibition of nearby PN dendrites.

Cortical SST-INs originate in the medial ganglionic eminence and migrate to their final destinations by postnatal day 5 (P5) in mice, but their connectivity and function are not yet fully mature ^20,21^. Recent work suggests global changes in the activity of SST-INs across cortical areas during the first postnatal week ^22^. Indeed, the intrinsic electrical properties of SST-INs in V1 and other cortical areas mature well after eye-opening at the end of the second postnatal week, leading to increased excitability by P28-29 ^23,24^. In turn, SST-INs may exert a critical influence on the maturation of cortical circuits, including the refinement of binocular receptive fields in V1 ^25^. In infragranular cortical circuits, SST-INs transiently receive thalamic input and regulate the maturation of parvalbumin-expressing interneurons^26^. Disruption of SST-IN activity has been implicated in neurodevelopmental disorders including autism ^27^ and schizophrenia ^28^ and loss of SST-INs during embryonic and early postnatal life leads to pathological neural activity and early death ^29^, further indicating a potentially critical role for these cells. Despite this evidence, the developmental trajectory of SST-IN activity and the early postnatal role of their synaptic inputs to PNs remains unknown.

Here, we examined the postnatal developmental trajectory of SST-INs in V1. Using 2-photon imaging to measure the visual and state-dependent responses of SST-INs and PNs in juvenile mice, we find that layer 2/3 V1 SST-INs gradually develop sensitivity to visual stimuli between P15 and P20. This emergence of visual responses arises in coordination with a substantial increase in excitatory synaptic innervation of SST-INs. SST-INs begin to exhibit size tuning and state-dependent modulation of spontaneous and visually evoked activity during this period, whereas nearby PNs exhibit visual feature selectivity and state-dependence from the earliest age studied. Targeted optogenetic manipulation of SST-INs in combination with 2-photon imaging of nearby PNs reveals that SST-INs have robust functional connectivity with local PNs by P15 but exert progressively more influence over the gain of visually evoked PN activity over time. Together, our results highlight the developmental emergence of a GABAergic cortical circuit mechanism for normalization that contributes to gain modulation of excitatory neuron responses.

## Results

### Emergence of visual sensitivity in SST-INs

In adult mice, SST-INs and PNs in layer 2/3 of V1 exhibit reliable, robust visual responses with a strong selectivity for large stimuli. Although extensive previous work has highlighted the early onset of well-tuned visual responses in PNs ^30,31^, relatively little is known about the development of visual sensitivity in SST-INs. We therefore used in vivo 2-photon imaging to directly test the developmental trajectories of visual sensitivity in both cell types following eye opening at P14. We imaged cellular activity in juvenile SST^Cre^;Ai148^F/0^ and Thy1-GCaMP6s mice constitutively expressing the calcium indicator GCaMP6 in SST-INs or PNs, respectively. We measured the activity of either SST-INs or PNs in head-fixed, awake behaving mice that ranged in age from P15, the first day after eye opening, to P29 (see Methods) (Fig. S1A, C). We observed no changes in the density of GCaMP-expressing SST-INs or the number of cells identified per imaging field of view in vivo for SST-INs or PNs, suggesting that GCaMP6 expression remained stable across ages (Fig. S1B, D-F).

Only ∼20% of SST-INs in awake behaving mice were responsive to visual stimulation at P15 (Fig. 1A-D). In contrast, by P21 over 80% of SST-INs were sensitive to visual stimuli, a proportion that was maintained at later ages (Fig. 1C-D). In good agreement with previous studies ^30,31^, we found a stable proportion of visually responsive PNs from P15 through P29 (Fig. 1E-F). Together, these data suggest distinct developmental timelines for visual sensitivity in PNs, whose receptive fields are well established prior to eye opening ^30,31^, and SST-INs, whose visual sensitivity increases rapidly starting at the end of the second postnatal week.

**Figure 1.**
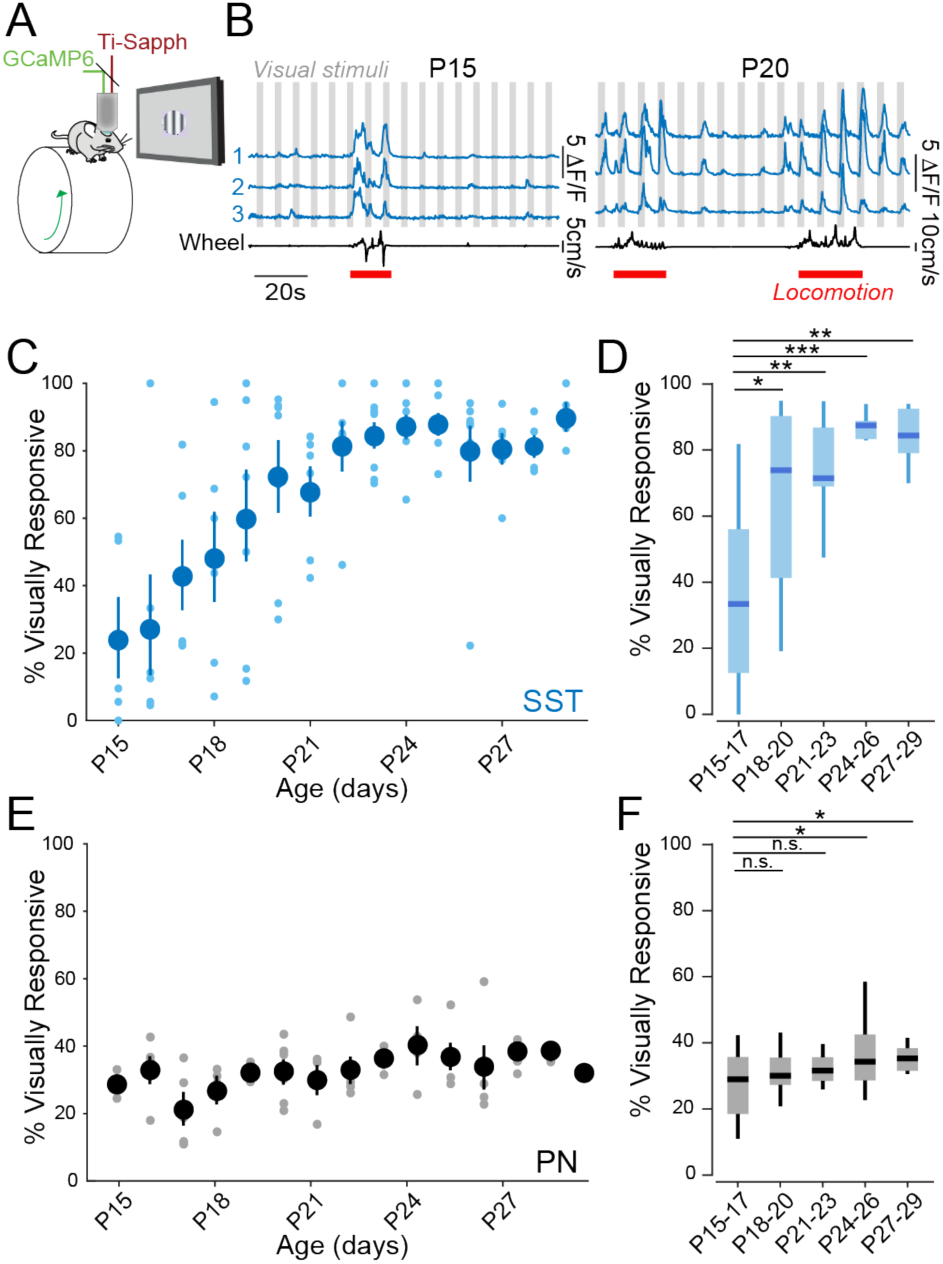
Visual sensitivity in SST-INs emerges following eye opening. (A) Schematic of the in vivo 2-photon imaging configuration. (B) Left: Ca2+ traces of three example P15 SST-INs (blue) recorded during the presentation of visual stimuli (gray) and wheel speed tracking (black) to identify locomotion bouts (red). Right: Ca2+ traces of three example P20 SST-INs. (C) Proportion of SST-INs that were visually responsive at each age. Large dark circles represent mean values and small light circles represent individual animals. Vertical lines show SEM. (D) Boxplots of the values in (C), aggregated into 3-day age groups (P15-17: n = 217 cells, 7 mice; P18-20: n = 184 cells, 8 mice; P21-23: n = 211 cells, 9 mice; P24-26: n = 230 cells, 10 mice; P27-29: n = 139 cells, 6 mice). Central mark indicates the median and whiskers indicate 25th and 75th percentiles. (E) and (F) Same as in (C) and (D) but for PNs (P15-17: n = 3301 cells, 6 mice; P18-20: n = 2791 cells, 7 mice; P21-23: n = 2425 cells, 6 mice; P24-26: n = 3638 cells, 6 mice; P27-29: n = 2099 cells, 6 mice). *p<0.05, **p<0.01, ***p<0.001, 0/1 inflated beta mixed-effects regression model with age as fixed effect and mouse as random effect.

Given the substantial increase in visual sensitivity in developing SST-INs, we examined whether excitatory innervation of these interneurons exhibited plasticity over the same period. We performed targeted whole-cell patch clamp recordings in acute slices from SST^Cre^;Ai9^F/0^ mice that constitutively expressed the red fluorescent indicator tdTomato selectively in SST-INs. We recorded miniature excitatory post-synaptic currents (mEPSCs) in SST-INs in layer 2/3 of V1 of mice ranging from P15 to P23. We found that the frequency of mEPSCs increased between P15 and P18 (Fig. 2A-C) whereas mEPSC amplitude (Fig. 2D) and 10%-90% rise time (Fig. 2E) remained unchanged. Together, these data suggest a rapid increase in excitatory synaptic innervation of SST-INs following eye opening, leading to emergence of visual responses by P21.

**Figure 2.**
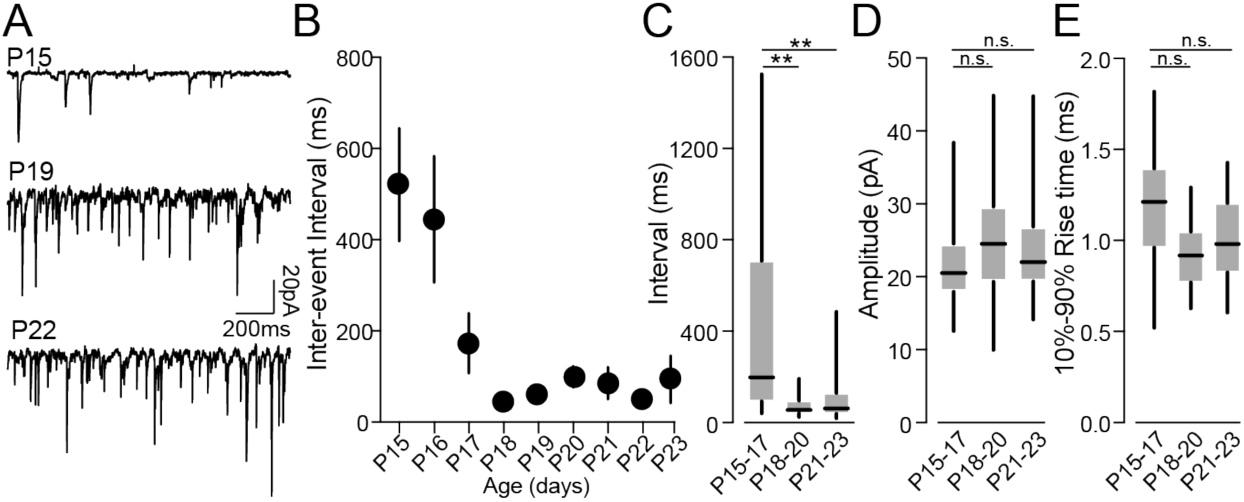
Rapid increase in excitatory synaptic input to SST-Ins. (A) Example traces of miniature EPSCs (mEPSCs) recorded ex vivo in SST-INs at P15 (upper), P19 (middle), and P22 (lower). (B) Inter-event intervals at each age P15-P23. Circles represent mean values and vertical lines show SEM. (C) Boxplots of the interval values in (B), aggregated into 3-day age groups. Central mark indicates the median and whiskers indicate 25th and 75th percentiles. (D) Boxplots of mEPSC amplitude across ages. (E) Boxplots of the 10%-90% rise time of mEPSCs across ages. P15-17: n = 30 cells. P18-20: n = 31 cells. P21-23: n = 20 cells. **p<0.01, One-way ANOVA with Tukey’s multiple comparisons test.

### Distinct developmental trajectories of stimulus selectivity in PNs and SST-INs

Previous work found that mature SST-INs and PNs in V1 exhibit robust, state-dependent feature selectivity for visual stimulus size ^8,13,16^. Indeed, SST-INs are highly sensitive to arousal and locomotion and are thought to potentially mediate surround suppression in nearby PNs and INs due to their responsiveness to large visual stimuli ^11,13,16^. However, little is known about the development of feature selectivity or state-dependent modulation of evoked activity in SST-INs. We found that neither SST-INs nor PNs exhibited overall population modulation of spontaneous activity levels by locomotion at P15-17, although individual cells in each population were positively or negatively modulated (Fig. S2A,B,D,F). By P18-21, SST-INs largely exhibited positive modulation by locomotion. In contrast, the PN population did not exhibit a change in state-dependent modulation across the P15-P29 age range (Fig. S2C, E-G).

At P15-17, SST-INs exhibited little visual sensitivity or modulation of visual responses by locomotion (Fig. 3A,B). Visual responses to stimuli of all sizes emerged by P18-20 and continued to increase in amplitude through P27-29 (Fig. 3B-D). SST-INs showing enhanced selectivity for large diameter stimuli by P21-23 (Fig. 3B, S3A-B), suggesting that progressive excitatory innervation drives changes in visual tuning over this period. In contrast, PNs exhibited robust visual responses that were selective for smaller stimuli by P15-17 and did not change significantly in amplitude between P15 and P29, further supporting a model where PN receptive fields and feature selectivity emerge prior to eye opening (Fig. 3B-D, S3C). The developmental trajectory of visual responses was similar across cells within each population regardless of visual tuning (Fig. S3D-E). SST-INs did not exhibit locomotion mediated gain modulation of their visual responses at P15-17 but developed robust modulation by P18-20 (Fig. 3E). Together, these data indicate that SST-INs develop robust visual responses and sensitivity to large stimuli well after PNs exhibit stable visual responses and sharp size tuning.

**Figure 3.**
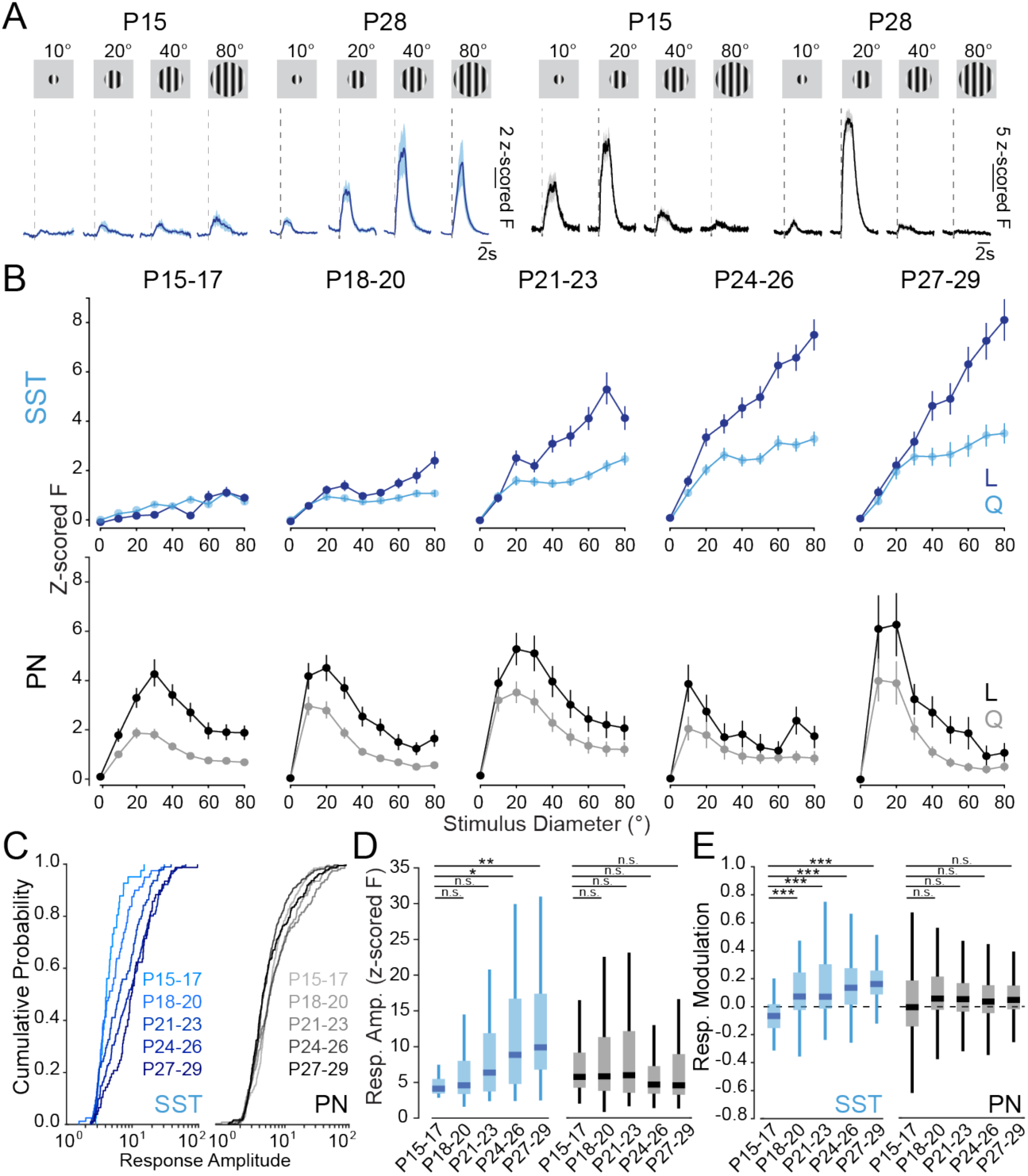
Developmental trajectory of SST-IN visual response amplitude and selectivity. (A) Responses of example SST-INs (blue) and PNs (black) to drifting grating stimuli of varying sizes at P15 and P28. Vertical dashed lines indicate visual stimulus onset. Shaded areas indicate mean ± SEM. (B) Population average visual responses of SST-INs (upper, blue) and PNs (lower, black) to stimuli of varying size across age groups. Responses are Z-scored to the 1-second baseline period before the stimulus onset for periods of quiescence (Q, light colors) and locomotion (L, dark colors). (C) Cumulative probability distribution of response amplitude at the preferred stimulus size for each age group of SST-INs (left; P15-17: n = 41 cells, 5 mice; P18-20: n = 80 cells, 7 mice; P21-23: n = 95 cells, 9 mice; P24-26: n = 141 cells, 10 mice; P27-29: n = 77 cells, 6 mice) and PNs (right; P15-17: n = 268 cells, 6 mice; P18-20: n = 217 cells, 7 mice; P21-23: n = 201 cells, 6 mice; P24-26: n = 397 cells, 6 mice; P27-29: n = 225 cells, 6 mice). (D) Boxplots of response amplitudes at the preferred stimulus size for each age group from (C) for SST-INs (blue) and PNs (black). Central mark indicates the median and whiskers indicate 25th and 75th percentiles. (E) Boxplots of locomotion-mediated gain modulation of visual response amplitudes in SST-INs (blue) and PNs (black) across ages. *p<0.05, **p<0.01, ***p<0.001, linear mixed-effects regression model with age as fixed effect and mouse as random effect.

### Functional impact of SST-INs on visual selectivity of PNs

To determine the influence of SST-INs on the local cortical circuit during postnatal development, we performed simultaneous 2-photon imaging and targeted optogenetic manipulations in juvenile mice. Using SST^Cre^; Thy1-GCaMP6 mice expressing a Cre-dependent ChrimsonR ^32^ construct (Fig. S4A, see Methods), we first selectively stimulated SST-INs while imaging the activity of local PNs at each age point. Activation of SST-INs during visual stimulation effectively silenced the visually-evoked activity of local PNs from P15 onwards (Fig. S4B-D), suggesting early establishment of functional synaptic connectivity between SST-INs and PNs despite minimal excitatory innervation of SST-INs.

We next examined the impact of SST-IN activity on the visual responses of local PNs by using optogenetic suppression of SST-IN activity via activation of a Cre-dependent ArchT construct ^33^ expressed in SST^Cre^; Thy1-GCaMP6 mice (Fig. 4A-B, S4E-G). We found that in juvenile PN cells that exhibited tuning for stimulus size, the impact of optogenetic suppression of SST-INs was diverse, with PNs showing either enhanced or reduced visual responses (Fig. 4C, S4H-I). In good agreement with our data on the visual responsiveness of SST-INs, we found that the impact of SST-IN suppression increased with age (P15-17 PNs: 8% enhanced, 36.2% reduced; P24-26 PNs: 20.2% enhanced, 30.1% reduced). In both cells whose visual responses were enhanced and reduced by suppressing SST-INs, the impact on visual tuning was strongest for stimuli ∼20 degrees, largely multiplicative at both early and late age points (Fig. 4D-E), and did not exhibit selective modulation of responses to large stimuli. Indeed, the impact of SST-IN suppression in reduced cells became more selective for smaller stimuli across ages (Fig. 4F, S4J). In contrast, the population impact of SST-IN suppression on PN spontaneous activity did not change across ages (Fig. S4K). Together, these data suggest that SST-INs largely exert a multiplicative gain modulation on PN visual responses at early and late ages, but their effect on the PN population becomes stronger across the postnatal period as SST-INs develop robust visually evoked activity.

**Figure 4.**
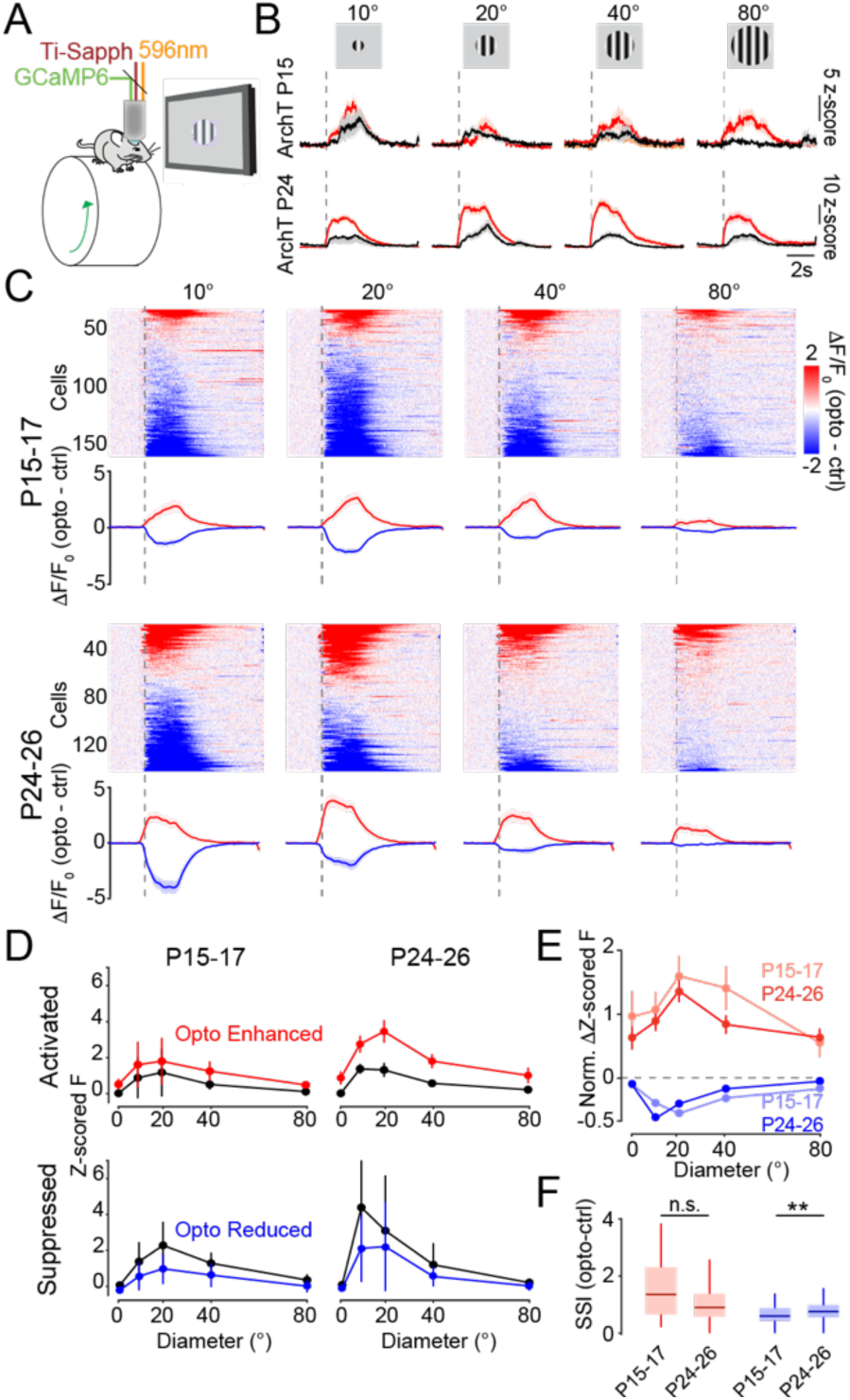
Emergence of SST-IN influence on visual selectivity in PNs. (A) Schematic of experimental configuration for simultaneous in vivo optogenetics and 2-photon imaging. (B) Example P15 and P24 PNs in SST^Cre+^;Thy1-GCaMP6 animals expressing Cre-dependent ArchT in SST-INs and GCaMP6 in PNs, showing visual responses to drifting grating stimuli of varying sizes during baseline conditions (black) and optogenetic suppression of SST-INs (red). Vertical dashed lines indicate visual stimulus onset. Shaded areas indicate mean ± SEM. (C) Subset of PNs exhibiting significant modulation of visually evoked responses by SST-IN suppression at varying stimulus sizes across ages. Each colored line represents a single PN’s activity, with either enhancement (red) or reduction (blue) of visual responses by the optogenetic stimulus. The difference in z-scored response between optogenetic and control trials for PNs showing enhanced (red) and reduced (blue) visual responses is plotted below each heat map. P15-17 (upper row; n = 229 cells, 6 mice), P24-26 (lower row; n = 206 cells, 4 mice). (D) Population average visual responses of PNs that were enhanced (upper row) and reduced (lower row) by SST-IN suppression across ages. Size tuning curves for enhanced (red) and reduced (blue) PNs are plotted against their control responses (black). (E). Normalized change in visual response amplitude at each stimulus size in enhanced (red) and reduced (blue) PNs at P15-17 (light colors) and P24-26 (dark colors). (F). Surround suppression index of the effect of optogenetic suppression of SST-INs for enhanced (red) and reduced (blue) PNs at each age. Enhanced PNs: P15-17 n = 28 cells, 6 mice; P24-26 n = 55 cells, 4 mice. Reduced PNs: P15 n = 123 cells, 6 mice; P24-26 n = 82 cells, 4 mice. **p<0.01, Mann-Whitney U test.

## Discussion

Our results reveal a key developmental window for the maturation of functional properties in the dendrite-targeting SST-IN population in cortical layer 2/3. We find that unlike PNs, in which visual receptive fields are established prior to eye opening, layer 2/3 V1 SST-INs have little response to visual stimuli at eye opening. Over the third postnatal week, SST-INs develop visual responses in coordination with a substantial increase in excitatory synaptic innervation. Over this same period, SST-IN responses exhibit increases in visual feature selectivity and modulation by changes in behavioral state. In contrast, PNs show robust visual responses and stable size tuning from P15 onwards. Finally, we find that SST-INs are functionally connected to PNs even at P15, but exert an increasing multiplicative impact on PN visual responses over the same time period.

SST-INs are among the earliest GABAergic populations to be integrated into the cortex, arriving in the cortical plate at P0 and concluding their laminar sorting by P5 ^21^. SST-INs in layers 4 and 5 undergo transient phases of connectivity early in postnatal life that are important for the maturation of thalamocortical and corticocortical circuits ^26,34–36^. In adult mice, layer 2/3 SST-INs receive extensive excitatory inputs which mostly arise from local, horizontally projecting PNs ^1,13^ and project to layer 1 cells and the dendrites of layer 2/3 PNs ^6,37,38^. The electrical properties of SST-INs in V1 and other cortical areas mature after eye-opening, with intrinsic excitability increasing until P28-29 ^23,24^ in contrast to PNs, which decline in intrinsic excitability ^39^. We found that the proportion of SST-INs exhibiting visually evoked responses was low at P15 and increased throughout the third postnatal week. Previous work has found that layer 2/3 PNs in V1 exhibit an increase in mEPSC rates but a decrease in mEPSC amplitudes during this period ^40^. In contrast, we found that the frequency of mEPSC events in SST-INs, but not their amplitude or rise time, rapidly increased after P15, indicating an increase in excitatory synaptic innervation and suggesting that developmental synaptic scaling may not occur similarly in all populations. However, the developmental trajectory of the synaptic and functional properties of GABAergic interneurons may vary across cortical areas ^41^. Indeed, previous findings in somatosensory (S1) cortex suggest that SST-INs in barrel cortex exhibit robust responses to whisker stimulation before P14 ^42^.

Mature SST-INs in V1 exhibit some selectivity for visual stimulus orientation and direction ^9,10,12^ and a unique responsiveness to large stimulus sizes ^8,13,16^. In juvenile animals at P15-17, we found relatively weak, untuned visual responses in these cells. In comparison, parvalbumin-expressing (PV) interneurons exhibit strong, selective visual responses by P17 and less tuning thereafter ^43,44^, suggesting distinct developmental trajectories in different GABAergic populations. These results also highlight potential differences in cortical development across species, as GABAergic interneurons in ferret V1 exhibit robust untuned responses to stimuli prior to eye opening and rapidly develop tuning following the onset of visual experience ^45^. Between P15 and P24, SST-INs developed stronger visual responses and adult-like preferences for large stimulus sizes. In good agreement with previous reports ^30,31^, we found that PNs at P15-17 already exhibited robust responses that were selective for small stimuli. As a result of their selectivity for large stimuli, mature SST-INs are thought to mediate surround suppression in PNs ^13^. However, the robust selectivity for small stimulus size in PNs at an age when SST-INs largely do not exhibit visual responses suggests that surround suppression is not regulated by SST-INs in this early postnatal period.

The activity of both excitatory and inhibitory neurons in cortical circuits is modulated by changes in behavioral state, such as arousal and locomotion ^12,16,46–51^. Mature PNs exhibit a broad distribution of responses to locomotion with a bias towards positive modulation, whereas SST-INs are largely positively modulated ^8,11,16^. We found that at the beginning of the third postnatal week, both PNs and SST-INs exhibited diverse responses to locomotion onset. SST-INs gradually developed positive state-dependent modulation over the following week. Locomotion also induces visual response gain modulation in diverse populations of mature V1 neurons, enhancing encoding of visual information during periods of arousal ^12,16,19,46–49,51^. Locomotion-induced gain modulation was initially absent in the SST-IN population at P15-17 and developed gradually through P27-29, suggesting that the developmental trajectories of visual responses and state modulation may be distinct. In contrast, PNs exhibited a more modest increase in visual gain modulation throughout this postnatal period. Previous work has implicated SST-INs in the mature cortex as a part of a VIP-SST-PN disinhibitory circuit, where state-dependent inhibition from VIP-INs may suppress SST activity and enhance PN activity ^8,16,50^. SST-INs are also directly and indirectly sensitive to cholinergic signals associated with locomotion and arousal ^52^. Together, these findings suggest a potential developmental window for state-dependent regulation of this circuit.

Synaptic connections between local SST-INs and PNs in layer 2/3 of V1 emerge by P7, with connection probability peaking at P11 and peak amplitude of inhibitory post-synaptic currents decreasing following eye opening ^20^. We found that activating SST-INs had a similar impact on PNs across the P15-P29 age range, supporting the idea that functional connectivity between these two cell types is established prior to eye opening. However, suppression of SST-IN activity during visual stimulation revealed a heterogenous ^53^ and age-dependent impact on PN visual responses. We did not observe an impact of SST-IN inhibition on the size tuning exhibited by PNs in the postnatal age range we examined, suggesting that SST-INs do not mediate surround suppression at these ages. Instead, SST-INs exerted an age-dependent increase in multiplicative modulation of PN responses, consistent with developmental emergence of an inhibitory circuit mechanism for normalization^54–56^.

Overall, our results highlight a window for the maturation of the functional properties of SST-INs in the postnatal cortex. We found a unique developmental trajectory for SST-INs in mouse primary visual cortex that was distinct from that of nearby excitatory pyramidal neurons, suggesting that the window between eye opening and the beginning of the classical critical period is a key time for the refinement of cortical operations that rely on dendrite-targeting interneurons. Overall, the maturation of functional properties in non-PV interneurons remains relatively poorly understood. However, the potential involvement of SST-INs in mechanisms of neurodevelopmental disorders including schizophrenia and autism ^27,28,57,58^ suggests that these cells may be critical mediators of cortical maturation and particularly vulnerable targets of postnatal dysregulation.

## Methods

### Animals

All animal handling and maintenance was performed according to the regulations of the Institutional Animal Care and Use Committee of the Yale University School of Medicine. Juvenile male and female Sst-IRES-Cre^+/+^(Jax stock no. 018973) crossed with Ai148^F/F^ (Ai148(TIT2L-GC6f-ICL-tTA2)-D, Jax stock no. 030328) (SST^Cre+^; Ai148^F/0^), Thy1-GCaMP6s (Jax stock no. 024275), and Sst-IRES-Cre^+/+^ crossed with Thy1-GCaMP6s (SST^Cre+^;Thy1-GCaMP6) mice were kept on a 12h light/dark cycle, provided with food and water ad libitum, and were returned to their parents and littermates following headpost implants. All mice used in the study were confirmed to have opened their eyes at P14. Juvenile mice were separated from their parents and housed by sex once they reached weaning age (P21). Imaging experiments were performed during the light phase of the cycle.

### Neonatal Local Injections

Local expression of the channelrhodopsin ChrimsonR or archaerhodopsin ArchT in V1 was achieved by intracranial injection. P0–P1 litters of Sst-IRES-Cre^+^;Thy1-GCaMP6s mice were removed from their home cage and placed on a heating pad. Pups were kept on ice for 8 min to induce anesthesia via hypothermia and then maintained on a metal plate surrounded by ice for the duration of the injection. Under a dissecting microscope, viral injections were made via beveled glass micropipette into the primary visual cortex (V1) at a depth of ∼350 um (QSI, Stoelting Co.). Pups were injected unilaterally with 1 μl of AAV9-hSyn-DIO-ChrimsonR-mRuby2-ST (2 × 10^12^ gc ml^−1^; Addgene #105448) or 1 μl of AAV9-CAG-Flex-ArchT (2 × 10^12^ gc ml^−1^; custom). Pups were then placed back on the heating pad with their littermates. Once the entire litter was injected, pups were gently rubbed with home cage bedding and nesting material and returned to their home cage.

### Headpost and Cranial Window Implantation Procedure

Implant surgeries were performed on juvenile mice (P15–P29), anesthetized with 1-2% isoflurane mixed with pure oxygen. Mice were required to weigh at least 6.0 grams at P15 to be considered for headpost implantation. During surgery, the scalp was first cleaned with Betadine solution. An incision was then made at the midline and the scalp resected to each side to leave an open area of the skull. After cleaning the skull and scoring it lightly with a surgical blade, a custom titanium head post was secured with C&B-Metabond (Butler Schein) with the left V1 centered. Skull screws were not used given the relative thinness of the mouse skull at these ages. A 3 mm^2^ craniotomy was made over the left V1. A glass window made of a 3 mm^2^ rectangular inner cover slip adhered with an ultraviolet-curing adhesive (Norland Products) to a 5 mm round outer cover slip (both #1, Warner Instruments) was inserted into the craniotomy and secured to the skull with Cyanoacrylate glue (Loctite). A circular ring was attached to the titanium headpost with glue, and additional Metabond was applied to cover any exposed skull. An analgesic (5 mg/kg Carprofen) and anti-inflammatory steroid (2 mg/mL Dexamethasone) was given immediately after surgery and on the two following days to aid recovery. Mice were given a course of antibiotics (Sulfatrim, Butler Schein) to prevent infection and returned to their littermates to provide maximum comfort for recovery.

### Histology

Following experiments, animals were given a lethal dose of sodium pentobarbital and perfused intracardially with 0.9% saline followed by cold 4% paraformaldehyde in 0.1 m sodium phosphate buffer. Brains were removed and fixed in 4% PFA/PBS solution for 24 hours and subsequently stored in PBS. Tissue was sectioned at 50mm using a vibrating blade microtome.

Widefield and confocal images were taken with a Zeiss LSM 900. To minimize counting bias we compared sections of equivalent bregma positions, defined according to the Mouse Brain atlas (Franklin and Paxinos, 2013). Cell counting was performed manually using a standardized 100 μm x 100 μm grid overlay to determine the average cell density in layers 2/3 of V1 across three consecutive sections.

### In Vivo Calcium Imaging and Optogenetic Stimulation

All imaging was performed during the second half of the light cycle in awake, behaving mice that were head-fixed so that they could freely run on a cylindrical wheel. A magnetic angle sensor (Digikey) attached to the wheel continuously monitored wheel motion. Mice were allowed to recover for at least an hour after window implantation before being fixed to the wheel and would comfortably run on the wheel within 3 consecutive days of imaging. The face (including the pupil and whiskers) was imaged with a miniature CMOS camera (Blackfly s-USB3, Flir) with a frame rate of 10 Hz.

Imaging was performed using a resonant scanner-based two-photon microscope (MOM, Sutter Instruments) coupled to a Ti:Sapphire laser (MaiTai DeepSee, Spectra Physics) tuned to 920 nm for GCaMP6. Emitted light was collected using a 25×1.05 NA objective (Olympus). Mice were placed on the wheel and head-fixed under the microscope objective. To prevent light contamination from the display monitor, the microscope was enclosed in blackout material that extended to the headpost. Images were acquired using ScanImage 4.2 at 30 Hz, 512×512 pixels. Imaging of layer 2/3 was performed at 150-350 μm depth relative to the brain surface. Each mouse was imaged for as many consecutive days as possible. For SST^Cre+^; Ai148^F/0^ mice, a single field of view could be imaged across consecutive days due to the relatively sparse distribution of the interneurons. In the Thy1-GCaMP6s mice, a different field of view was imaged for each consecutive day of imaging. Visual stimulation, wheel position, and Ca2+ imaging microscope resonant scanner frame ticks were digitized (5 kHz) and collected through a Power 1401 (CED) acquisition board using Spike 2 software.

Optogenetic stimulation was achieved by aligning a 594nm laser to the same light path as the Ti:Sapphire laser of the two-photon microscope, allowing us to activate ChrimsonR or ArchT without affecting imaging quality. The main dichroic of the microscope was replaced with one with a 594nm notch to allow dual IR and 594nm excitation. Optogenetic stimulation was delivered through the two-photon microscope objective lens as a single pulse of light beginning 250 ms prior to the onset of the visual stimulus and concluding at the end of the 2 second visual stimulus period. This stimulation was delivered during alternating visual stimuli.

### Visual Stimulation

Visual stimuli were generated using Psychtoolbox-3 in MATLAB and presented on a gamma-calibrated LCD monitor (17 inches) at a spatial resolution of 1280 x 960, a real-time frame rate of 60Hz, and a mean luminance of 30 cd/m^2^ positioned 20 cm from the right eye. Stimuli had a temporal frequency of 2 Hz, spatial frequency of 0.04 cycles per degree, and orientation of 180°. To center stimuli on the receptive field, 100% contrast stimuli were randomly presented in nine 3×3 sub-regions to identify the location that evoked the largest population response in the field of view. The screen was centered, and the process was repeated until a center was identified. Stimuli in each session were randomized and presented in blocks with a fixed duration of 2 s and an interstimulus interval of 5 s, with a mean-luminance gray screen between stimuli. For size tuning, the visual angle was linearly spaced from 0 to 80° in diameter in steps of 10°, where each size was presented 45 times. For the optogenetics experiments, the sizes presented were limited to 0, 10, 20, 40, and 80° in diameter in order to accommodate a sufficient number of optogenetic and control trials for statistical comparison.

### Ex vivo electrophysiology

Under isoflurane anesthesia, mice from each age group were decapitated and transcardially perfused with ice-cold choline-artificial cerebrospinal fluid (choline-ACSF) containing (in mM): 110 choline, 25 NaHCO_3_, 1.25 NaH_2_PO_4_, 2.5 KCl, 7 MgCl_2_, 0.5 CaCl_2_, 20 glucose, 11.6 sodium ascorbate, 3.1 sodium pyruvate. Acute coromal slices (300 μm) were prepared from the left hemisphere and transferred to ACSF solution containing (in mM): 127 NaCl, 25 NaHCO_3_, 1.25 NaH_2_PO_4_, 2.5 KCl, 1 MgCl_2_, 2 CaCl_2_, and 20 glucose bubbled with 95% O_2_ and 5% CO_2_. After an incubation period of 30 min at 32°C, the slices were maintained at room temperature until use. Visualized whole-cell recordings were performed by targeting fluorescently labeled SST-INs in the primary visual cortex (V1). All recordings were performed at room temperature. Series resistance (Rs) values were <20 MΩ and uncompensated. For miniature excitatory postsynaptic current recordings (mEPSC), the ACSF contained 1 µM TTX to block sodium channels and 10μM gabazine to block GABAergic currents. The internal solution contained (in mM): 126 cesium gluconate, 10 HEPES, 10 sodium phosphocreatine, 4 MgCl_2_, 4 Na_2_ATP, 0.4 Na_2_GTP, 1 EGTA (pH 7.3 with CsOH). Cells were voltage-clamped at –70 mV. For miniature postsynaptic current recordings, the ACSF contained 1 uM TTX to block sodium channels. For mEPSCs, the internal solution contained (in mM): 126 cesium gluconate, 10 HEPES, 10 sodium phosphocreatine, 4 MgCl2, 4 Na2ATP, 0.4 Na2GTP, 1 EGTA (pH 7.3 with CsOH).

## Data analysis

### Wheel Position and Changepoints

Wheel position was determined from the output of the linear angle detector. The circular wheel position variable was first transformed to the [-π, π] interval. The phases were then circularly unwrapped to get running distance as a linear variable, and locomotion speed was computed as a differential of distance (cm/s). A change-point detection algorithm detected locomotion onset/offset times based on changes in standard deviation of speed. Locomotion onset or offset times were estimated from periods when the moving standard deviations, as determined in a 0.5s window, exceeded or fell below an empirical threshold of 0.1. Locomotion trials were required to have average speed exceeding 0.25 cm/s and last longer than 1 s. Quiescence trials were required to last longer than 2 s and have an average speed < 0.25 cm/s.

### Quantification of Calcium Signals

Analysis of imaging data was performed using ImageJ and custom routines in MATLAB (The Mathworks). Motion artifacts and drifts in the Ca2+ signal were corrected with the moco plug-in in ImageJ (Dubbs et al., 2016), and regions of interest (ROIs) were selected as previously described (Chen et al., 2013). All pixels in each ROI were averaged as a measure of fluorescence, and the neuropil signal was subtracted (Chen et al, 2013; Lur et al., 2016; Tang et al., 2020). Excitatory pyramidal cell data was processed through Suite2p (Pachitariu, et al. 2017) due to the high prevalence of these cells in the fields of view imaged but the output was similarly analyzed in the same custom routines in MATLAB.

### Modulation Index

For modulation by behavioral state without visual stimulation, we used the spontaneous periods recorded as described above and selected locomotion trials that lasted 5 s or longer and quiescent trials that lasted 20 s or longer. To determine whether Ca2+ activity was altered during behavioral state transitions, ΔF/F(t) from [0,5]s after locomotion onset (Ca_L-ON_) was compared with ΔF/F(t) from [10,15]s after locomotion offset (Ca_Q_) by computing a modulation index (MI), where MI = (Ca_L-ON_ – Ca_Q_)/(Ca_L-ON_ +Ca_Q_). A minimum of 5s of quiescence after this period [15,20]s was required to prevent anticipatory effects on Ca_Q_. To ascertain the significance of this MI, we used a shuffling method in which the wheel trace was randomly circularly shifted relative to the fluorescence trace 1,000 times. Cells were deemed significantly modulated if their MI was outside of the 95% confidence interval of the shuffled comparison.

### Visual Responses

Visual response amplitude was determined as the peak of the z-scored change in fluorescence during the 2s visual stimulus (F) compared to the 1s baseline before the stimulus (F_0_), given by (F-μ_F0_)/α_F0_. To reduce high-frequency noise when selecting peak amplitude, we applied MATLAB’s zero-phase filtering (*filtfilt*) using a second-order infinite impulse response low-pass filter with a half-power frequency of 3 Hz. When separating the effects of state, the mouse was required to be running (or sitting) during the majority (>50%) of each of the 1s baseline and the 2s visual stimulation. To evaluate visual responsiveness, we conducted a t-test comparing peak response amplitudes to the preferred stimulus size against responses to a blank stimulus (zero degrees). The preferred stimulus size was determined as the size yielding the highest mean response amplitude across all tested sizes. We identified suppressed cells using bootstrap analysis with 1000 resamplings to compute the median response amplitudes from the z-scored fluorescence traces. Cells with a median bootstrap amplitude below zero were classified as suppressed and excluded from the positively visually responsive group.

Size tuning of all cell types, and particularly of SST-INs, prefers larger stimulus sizes when not well centered (Dipoppa et al., 2018). Tuned cells were identified using a t-test comparing the responses to their preferred stimulus size with those to the largest size. Cells with a statistically significant difference (α = 0.05) in peak responses were classified as tuned, and those without a significant difference as untuned.

### Optogenetic Responses

We assessed whether a cell was significantly modulated by optogenetic stimulation by comparing peak response to the preferred stimulus size during optogenetic versus control trials using the Mann-Whitney U test (α = 0.05). Modified cells were categorized as enhanced or reduced based on whether their response amplitude increased or decreased, respectively, during optogenetic trials compared to control. Optogenetic modulation was quantified after normalizing the data to the peak response of the control trials for each cell. This normalization facilitated comparison across cells with varying response amplitudes. The amount of modulation was determined by calculating the difference in normalized peak responses between optogenetic and control trials, providing a measure of the optogenetic effect’s magnitude and direction. To determine how the response varies over stimulus sizes, we calculated suppression indices for both control and optogenetic trials by determining the difference between the response to the preferred stimulus size and the response to the largest size. A change in suppression index was quantified as the difference between the optogenetic and control indices.

### Quantification and Statistical Analyses

When possible, we used mixed effect regression models for imaging data, due to its nested structure with multiple cells recorded within each mouse. We treated the age group as the fixed effect, while individuals (mice) were random effects. For the PN control data, which uniquely included multiple fields of view per mouse, fields of view were additionally modeled as nested random effects within mice to capture within-subject variability. Model complexity, including the addition of nested random effects, was evaluated based on Akaike and Bayesian Information Criteria (AIC/BIC). We additionally evaluated the fit of our models by checking homoscedasticity in plots of residuals vs. fitted values, Q-Q plots to assess residual normality, and an analysis of deviance residuals to detect overdispersion and model fit anomalies.

For response variables that were continuous and normally distributed, we used a linear mixed effect model, implemented with the *lmer* function from the *lme4* package in R (Version 4.2.2), R Core Team (2022). For continuous but bounded variables, such as the percent of visually responsive cells, we instead used a 0/1 inflated beta mixed effect regression model. For these data, the variables were scaled to [0, 1] to represent the proportion compared to maximum. Then we fit a 0/1 inflated beta mixed effect regression model using the *gamlss* package in R using the family “BEINF” (R Version 4.2.2), R Core Team (2022). After fitting the model, we transformed the estimates back to probabilities using the inverse link function to model coefficients to facilitate their interpretation.

To compare the number of SST-INs per field of view (counts), we used a Poisson mixed effects model, with age as the fixed effect and the individuals (mice) as the random effect. The PN data was overdispersed (the variance was significantly larger than the mean). To address overdispersion in our data, we evaluated both Poisson and negative binomial distributions for our mixed effects modeling. The models were fitted using the Poisson family from *glmer* and *glmer.nb* for the negative binomial model (*lme4* package) in R. Model selection was based on the likelihood ration test (*lrtest* from *lmtest* package in R, Version 4.2.2), R Core Team (2022). The negative binomial model was chosen due to significant improvement in model fit (p < 0.05).

To assess differences among age groups, post-hoc analyses were conducted using the *emmeans* package in R, employing the Dunnett correction method for multiple comparisons. This method facilitated the comparison of each older age group to the youngest age group (P15-17) as a reference, effectively controlling for the family-wise error rate in the context of multiple testing.

For one subset of cells, the mixed effect models did not converge. In this circumstance, we were comparing visually responsive and tuned PNs, that were either positively or negatively modulated by the suppression of SST-INs, across two age groups (P15-17 and P24-26). We instead employed the Mann-Whitney U test (after a Shapiro-Wilk test that showed the data did not follow a normal distribution).

For the linear mixed effect model, the response variable was modeled as: response ∼ age + (1 | mouse), which has the following mathematical form:

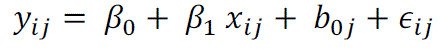

where *y_ijk_* is the *i^th^* observation for the *j^th^* mouse. *x_ij_* is the age group for the observation *i* of the *j^th^* mouse. β_0_ is the intercept, β_1_ is the effect of age, *b*_0,1_ is the random intercept for the *j^th^*mouse and captures the deviation of the *j^th^* mouse’s baseline level from the overall intercept β_0_, and ϵ*_ijk_* is the residual error for the i^th^ observation within the j^th^ mouse. The random effects have prior distributions 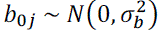, and the error term has the distribution 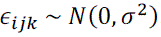.

All of the details of the statistical tests used, including n’s and definition of center and dispersion, are provided in Table 1. No tests were used to justify sample size, but sample sizes in the current study are comparable to several recent studies in behaving mice ^8,17,51,59^.

**Table 1.** Summary of all statistical analyses.

| Figure | Comparison | N | Test | Estimate | Test statistic | 95% Confidence Interval | p-value |
| --- | --- | --- | --- | --- | --- | --- | --- |
| 1D | Comparison of percent visually responsive cells for SST between P15-17 and each other age group | P15-17: n = 217 cells, 7 mice; P18-20: n = 184 cells, 8 mice; P21-23: n = 211 cells, 9 mice; P24-26: n = 230 cells, 10 mice; P27-29: n = 139 cells, 6 mice | Zero/one inflated beta mixed effects regression model (Age as fixed effect; mouse as random effect). Log-odds estimates transformed to probabilities. P-values corrected for multiple comparisons via EMMMeans (Dunnett) | 0.76621 | 2.67333 | 0.52443 to 0.90689 | P15-17 vs. P18-20: 0.02729 |
|  |  |  |  | 0.81339 | 3.35193 | 0.59744 to 0.92754 | P15-17 vs. P21-23: 0.00308 |
|  |  |  |  | 0.83917 | 3.71362 | 0.63664 to 0.93954 | P15-17 vs. P24-26: 0.00080 |
|  |  |  |  | 0.85669 | 3.57164 | 0.63645 to 0.95330 | P15-17 vs. P27-29: 0.00138 |
| 1F | Comparison of percent visually responsive cells for PYR between P15-17 and each other age group | P15-17: n = 3301 cells, 6 mice; P18-20: n = 2791 cells, 7 mice; P21-23: n = 2425 cells, 6 mice; P24-26: n = 3638 cells, 6 mice; P27-29: n = 2099 cells, 6 mice | Zero/one inflated beta mixed effects regression model (Age as fixed effect; mouse as random effect). Log-odds estimates transformed to probabilities. P-values corrected for multiple comparisons via EMMMeans (Dunnett) | 0.54329 | 1.11808 | 0.44837 to 0.63516 | P15-17 vs. P18-20: 0.60821 |
|  |  |  |  | 0.56802 | 1.74350 | 0.47218 to 0.65903 | P15-17 vs. P21-23: 0.24513 |
|  |  |  |  | 0.60887 | 2.95497 | 0.51879 to 0.69211 | P15-17 vs. P24-26: 0.01169 |
|  |  |  |  | 0.60447 | 2.72930 | 0.51073 to 0.69111 | P15-17 vs. P27-29: 0.02321 |
| S1B | Comparison of number of SST-INs per field of view between P15-17 and each other age group | P15-17: n = 7 mice; P18-20: n = 8 mice; P21-23: n = 9 mice; P24-26: n = 10 mice; P27-29: n = 6 mice | Poisson mixed effects regression model (Age as fixed effect; mouse as random effect). Estimates transformed to linear scale. P-values corrected for multiple comparisons via EMMMeans (Dunnett) | 0.94920 | -0.47414 | 0.72481 to 1.24307 | P15-17 vs. P18-20: 0.93934 |
|  |  |  |  | 1.03486 | 0.27753 | 0.76443 to 1.40096 | P15-17 vs. P21-23: 0.98290 |
|  |  |  |  | 1.14088 | 1.01937 | 0.83080 to 1.56671 | P15-17 vs. P24-26: 0.67135 |
|  |  |  |  | 1.09309 | 0.62744 | 0.77183 to 1.54807 | P15-17 vs. P27-29: 0.88403 |
| S1D | Comparison of number of PN's per field of view between P15-17 and each other age group | P15-17: n = 6 mice; P18-20: n = 7 mice; P21-23: n = 6 mice; P24-26: n = 6 mice; P27-29: n = 6 mice | Negative Binomial mixed effects regression model (Age as fixed effect; mouse as random effect). Estimates transformed to linear scale. P-values corrected for multiple comparisons via EMMMeans (Dunnett) | 0.79027 | -1.55473 | 0.54511 to 1.14568 | P15-17 vs. P18-20: 0.33931 |
|  |  |  |  | 0.74632 | -1.81296 | 0.50233 to 1.10883 | P15-17 vs. P21-23: 0.21518 |
|  |  |  |  | 1.06438 | 0.37145 | 0.70494 to 1.60709 | P15-17 vs. P24-26: 0.96586 |
|  |  |  |  | 0.72617 | -1.83272 | 0.47320 to 1.11438 | P15-17 vs. P27-29: 0.20713 |
| S1F | Comparison of density of layer 2/3 SST-INs between P15-17 and P27-29 | P15-17: n = 4 mice; P27-29: n = 4 mice | Mann-Whitney U test |  | U = 6 |  | 0.6857 |
| 2C | Comparison of mEPSC | P15-17: 30 cells<br>P18-20: 31 cells |  |  |  | 179.7 to 531.3 | <0.0001 |
|  | interval for SST between P15-17 and each other age group | P21-23: 20 cells | One-way ANOVA; Tukey's multiple comparisons test |  |  | 128.3 to 515.5 | 0.0005 |
| 2D | Comparison of SST mEPSC amplitude between P15-17 and each other age group | P15-17: 30 cells<br>P18-20: 31 cells<br>P21-23: 20 cells | One-way ANOVA; Tukey's multiple comparisons test |  |  | -6.917 to 1.992 | 0.3879 |
|  |  |  |  |  |  | -6.607 to 3.435 | 0.7317 |
| 2E | Comparison of the 10-90% rise time for SST mEPSCs between P15-17 and each other age group | P15-17: 30 cells<br>P18-20: 31 cells<br>P21-23: 20 cells | One-way ANOVA; Tukey's multiple comparisons test |  |  | -0.5549 to 1.088 | 0.7191 |
|  |  |  |  |  |  | -0.7261 to 1.078 | 0.8871 |
| S2G left | Comparison of locomotion modulation indices for SST between P15-17 and each other age group | P15-17: n = 114 cells, 4 mice;<br>P18-20: n = 180 cells, 5 mice;<br>P21-23: n = 211 cells, 5 mice;<br>P24-26: n = 246 cells, 8 mice;<br>P27-29: n = 142 cells, 3 mice | Linear mixed effects regression model (Age as fixed effect; mouse as random effect). P-values corrected for multiple comparisons via EMMeans (Dunnett) | 0.35966 | 9.63621 | 0.26791 to 0.45142 | P15-17 vs. P18-20: 0.00000 |
|  |  |  |  | 0.17755 | 4.15703 | 0.07240 to 0.28270 | P15-17 vs. P21-23: 0.00015 |
|  |  |  |  | 0.35551 | 9.64861 | 0.26488 to 0.44614 | P15-17 vs. P24-26: 3.54298E-10 |
|  |  |  |  | 0.27307 | 3.68158 | 0.08471 to 0.46144 | P15-17 vs. P27-29: 0.00224 |
| S2G right | Comparison of locomotion modulation indices for PYR between P15-17 and each other age group | P15-17: n = 3301 cells, 6 mice, 13 fov; P18-20: n = 2545 cells, 7 mice, 12 fov; P21-23: n = 2190 cells, 6 mice, 11 fov; P24-26: n = 3638 cells, 6 mice, 14 fov; P27-29: n = 2267 cells, 6 mice, 12 fov | Linear mixed effects regression model (Age as fixed effect; mouse with nested field of view (fov) as random effect). P-values corrected for multiple comparisons via EMMeans (Dunnett) | 0.01486 | 1.39615 | -0.01206 to 0.04178 | P15-17 vs. P18-20: 0.43954 |
|  |  |  |  | 0.02731 | 2.21614 | -0.00389 to 0.05852 | P15-17 vs. P21-23: 0.10271 |
|  |  |  |  | 0.01907 | 1.46043 | -0.01462 to 0.05275 | P15-17 vs. P24-26: 0.40773 |
|  |  |  |  | 0.02294 | 1.73302 | -0.01103 to 0.05691 | P15-17 vs. P27-29: 0.26733 |
| 3D left | Comparison of response amplitude to preferred stimulus size for SST between P15-17 and each other age group | P15-17: n = 41 cells, 5 mice; P18-20: n = 80 cells, 7 mice; P21-23: n = 95 cells, 9 mice; P24-26: n = 141 cells, 10 mice; P27-29: n = 77 cells, 6 mice | Linear mixed effects regression model (Age as fixed effect; mouse as random effect). P-values corrected for multiple comparisons via EMMeans (Dunnett) | 1.06526 | 0.47960 | -4.42811 to 6.55863 | P15-17 vs. P18-20: 0.93770 |
|  |  |  |  | 1.08844 | 0.49445 | -4.36249 to 6.53937 | P15-17 vs. P21-23: 0.93311 |
|  |  |  |  | 5.53277 | 2.60764 | 0.27445 to 10.7911 | P15-17 vs. P24-26: 0.03565 |
|  |  |  |  | 7.84966 | 3.32058 | 1.99234 to 13.7070 | P15-17 vs. P27-29: 0.00421 |
| 3D right | Comparison of response amplitude to preferred stimulus size for PYR between P15-17 and | P15-17: n = 268 cells, 6 mice, 14 fov; P18-20: n = 217 cells, 7 mice, 13 fov; P21-23: n = 201 cells, 6 mice, 12 fov; P24-26: n = 397 cells, 6 mice, 14 | Linear mixed effects regression model (Age as fixed effect; mouse with nested field of view (fov) as random effect). P-values corrected for multiple comparisons via | 2.14935 | 1.51876 | -1.43791 to 5.73662 | P15-17 vs. P18-20: 0.36989 |
|  |  |  |  | 3.71669 | 2.49685 | -0.09191 to 7.52529 | P15-17 vs. P21-23: 0.05766 |
|  |  |  |  | -0.32900 | -0.23611 | -4.01056 to 3.35256 | P15-17 vs. P24-26: 0.98828 |
|  | each other age group | fov; P27-29: n = 225 cells, 6 mice, 12 fov | EMMeans (Dunnett) | 0.75607 | 0.51393 | -3.06280 to 4.57494 | P15-17 vs. P27-29: 0.92694 |
| 3E left | Comparison of visual response gain modulation for SST between P15-17 and each other age group | P15-17: n = 48 cells, 5 mice; P18-20: n = 106 cells, 8 mice; P21-23: n = 130 cells, 9 mice; P24-26: n = 172 cells, 10 mice; P27-29: n = 110 cells, 6 mice | Linear mixed effects regression model (Age as fixed effect; mouse as random effect). P-values corrected for multiple comparisons via EMMeans (Dunnett) | 0.29579 | 8.05251 | .20537 to 0.38621 | P15-17 vs. P18-20: 2.04341E-11 |
|  |  |  |  | 0.31608 | 8.56916 | 0.22525 to 0.40691 | P15-17 vs. P21-23: 2.22045E-16 |
|  |  |  |  | 0.34631 | 9.48657 | 0.25640 to 0.43622 | P15-17 vs. P24-26: 0.00000 |
|  |  |  |  | 0.35354 | 8.75164 | 0.25405 to 0.45304 | P15-17 vs. P27-29: 0.00000 |
| 3E right | Comparison of visual response gain modulation for PYR between P15-17 and each other age group | P15-17: n = 692 cells, 6 mice, 12 fov; P18-20: n = 635 cells, 7 mice, 13 fov; P21-23: n = 542 cells, 6 mice, 12 fov; P24-26: n = 992 cells, 6 mice, 14 fov; P27-29: n = 558 cells, 6 mice, 12 fov | Linear mixed effects regression model (Age as fixed effect; mouse with nested field of view (fov) as random effect). P-values corrected for multiple comparisons via EMMeans (Dunnett) | 0.02020 | 0.78415 | -0.04481 to 0.08520 | P15-17 vs. P18-20: 0.81039 |
|  |  |  |  | -0.02072 | -0.71961 | -0.09348 to 0.05204 | P15-17 vs. P21-23: 0.84287 |
|  |  |  |  | -0.02288 | -0.72786 | -0.10358 to 0.05783 | P15-17 vs. P24-26: 0.83928 |
|  |  |  |  | -0.01036 | -0.32593 | 0.07087 to -0.32593 | P15-17 vs. P27-29: 0.97490 |
| 4F left | Comparison of SSI in enhanced PNs in response to SST-IN suppression between P15-17 and each other age group | P15-17: n = 28 cells, 6 mice, P24-26: n = 55 cells, 4 mice | Mann-Whitney U test |  | 1.76737 |  | P15-17 vs. P24-26: 0.07717 |
| 4F right | Comparison of SSI in reduced PNs in response to SST-IN suppression between P15-17 and each other age group | P15-17: n = 123 cells, 6 mice, P24-26: n = 82 cells, 4 mice | Mann-Whitney U test |  | -3.19277 |  | P15-17 vs. P24-26: 0.00141 |
| S4D | Comparison of PN response to activation of SST-INs between each age group and zero | P15-17: n = 124 cells, 6 mice; P18-20: n = 268 cells, 7 mice; P21-23: n = 133 cells, 3 mice; P24-26: n = 264 cells, 6 mice; P27-29: n = 75 cells, 1 mouse | Intercept-only linear mixed effects regression model (No fixed effect; mouse as random effect). P-values corrected for multiple comparisons (Benjamini & Hochberg) | -3.24817 | -9.95641 | -3.89394 to -2.60240 | P15-17 vs. zero: 2.60617E-17 |
|  |  |  |  | -5.44532 | -14.9913 | -6.16048 to -4.73016 | P18-20 vs. zero: 1.68733E-36 |
|  |  |  |  | -2.72252 | -10.6309 | -3.22910 to -2.21594 | P21-23 vs. zero: 3.77691E-19 |
|  |  |  |  | -3.85674 | -11.2906 | -4.52933 to -3.18415 | P24-26 vs. zero: 6.87762E-24 |
|  |  |  |  | -2.88503 | -6.00191 | -3.84281 to -1.92724 | P27-29 vs. zero: 8.00089E-08 |
| S4K left | Comparison of change in spontaneous activity for enhanced PNs between P15-17 and P24-26 | P15-17: n = 202 cells, 6 mice; P24-26: n = 193 cells, 4 mice. | Linear mixed effects regression model (Age as fixed effect; mouse as random effect) | -0.01139 | -0.14107 | -0.17013 to 0.14735 | 0.88788 |
| S4K<br>right | Comparison<br>of change in<br>spontaneous<br>activity for<br>reduced PN<br>between<br>P15-17 and<br>P24-26 | P15-17: n = 166<br>cells, 6 mice;<br>P24-26: n = 124<br>cells, 4 mice. | Linear mixed effects<br>regression model<br>(Age as fixed effect;<br>mouse as random<br>effect) | 0.01795 | 0.54215 | -0.04720 to 0.08309 | 0.58813 |

## Acknowledgements

The authors thank all members of the Higley and Cardin laboratories for helpful input throughout all stages of this study. We thank Rima Pant for generation of AAV vectors and Lauren Panzera for consulting on cloning. This work was supported by funding from the NIH (R01EY022951 to JAC, R01EY035127 to JAC, R01MH113852 to JAC and MJH, R01MH099045, R21MH121841, and DP1EY033975 to MJH, F31EY032793 to AW, K99EY030549 to KAF, EY026878 to the Yale Vision Core, T32 EY022312 support for AW), and a BBRF Young Investigator Grant (to KAF).

## Author Contributions

AW, KAF, MJH, and JAC designed the experiments. AW and JG collected the data. KAF analyzed the in vivo data and JG analyzed the ex vivo data. AW, KAF, and JAC wrote the manuscript.

## Inclusion and diversity

We support inclusive, diverse, and equitable conduct of research.

---

**Supplemental Figure 1.**
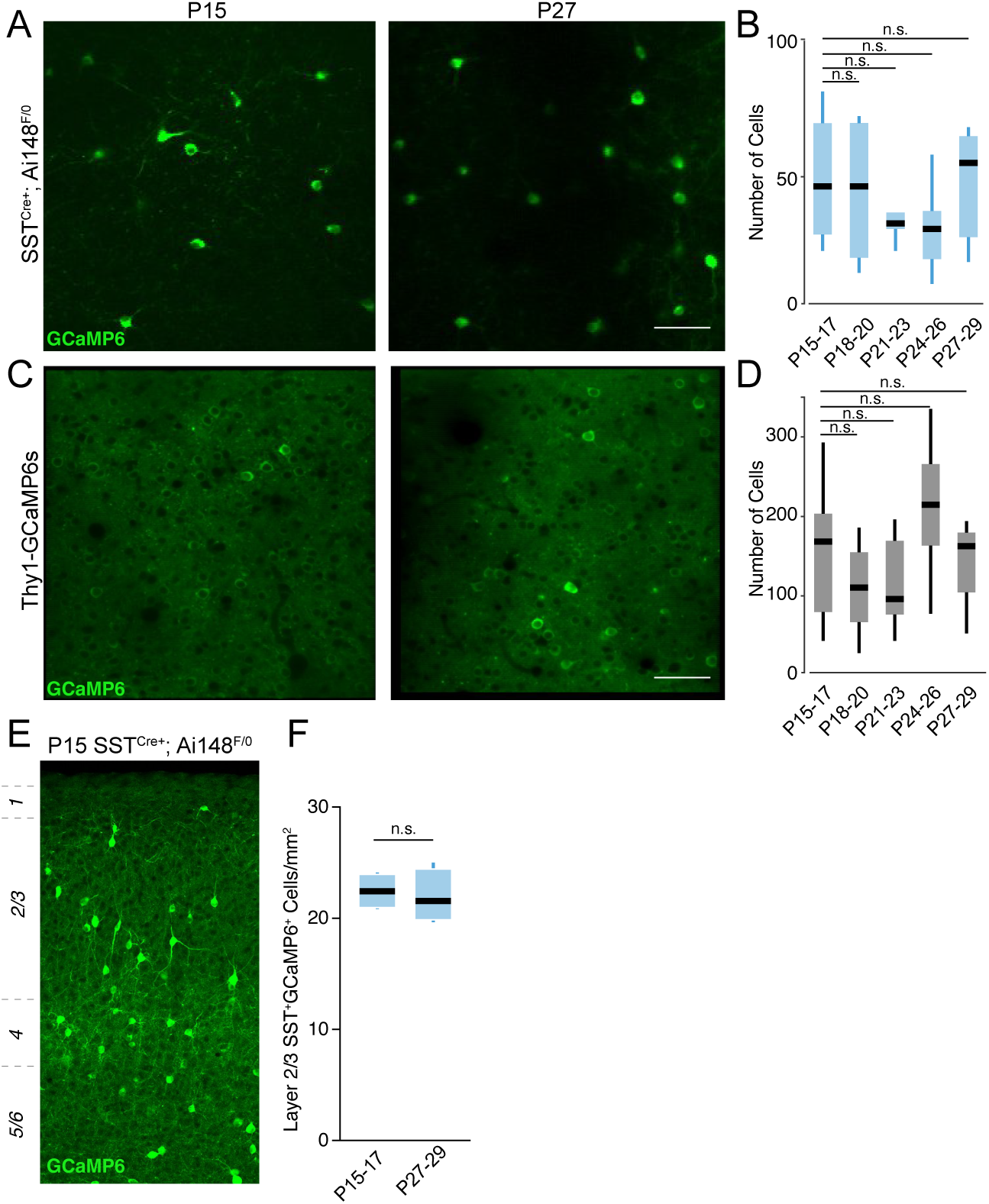
In vivo 2-photon imaging of GCaMP6-expressing SST-INs and PNs (green) across ages. (A) Example in vivo fields of view of SST-INs at P15 (left) and P27 (right). (B) Population average number of SST-INs recorded per field of view in each age group. P15-17: n = 7 mice; P18-20: n = 8 mice; P21-23: n = 9 mice; P27-29: n = 6 mice. Poisson mixed effects regression model with age as fixed effect and mouse as random effect. (C and D) Same as in (A) and (B) but for PNs. P15-17: n = 6 mice; P18-20: n = 7 mice; P21-23: n = 6 mice: P24-26: n = 6 mice; P27-29: n = 6 mice. Negative binomial mixed effects regression model with age as fixed effect and mouse as random effect. Scale bars denote 50μm. E. Example section from a P15 SST^Cre+^;Ai148^F/0^ mouse showing GCaMP6-expressing SST-INs (green) throughout the cortex. F. Population average density of SST-INs in cortical layer 2/3 at P15 and P27. P15-17: n = 4 mice; P27-29: n = 4 mice. Mann-Whitney U test.

**Supplemental Figure 2.**
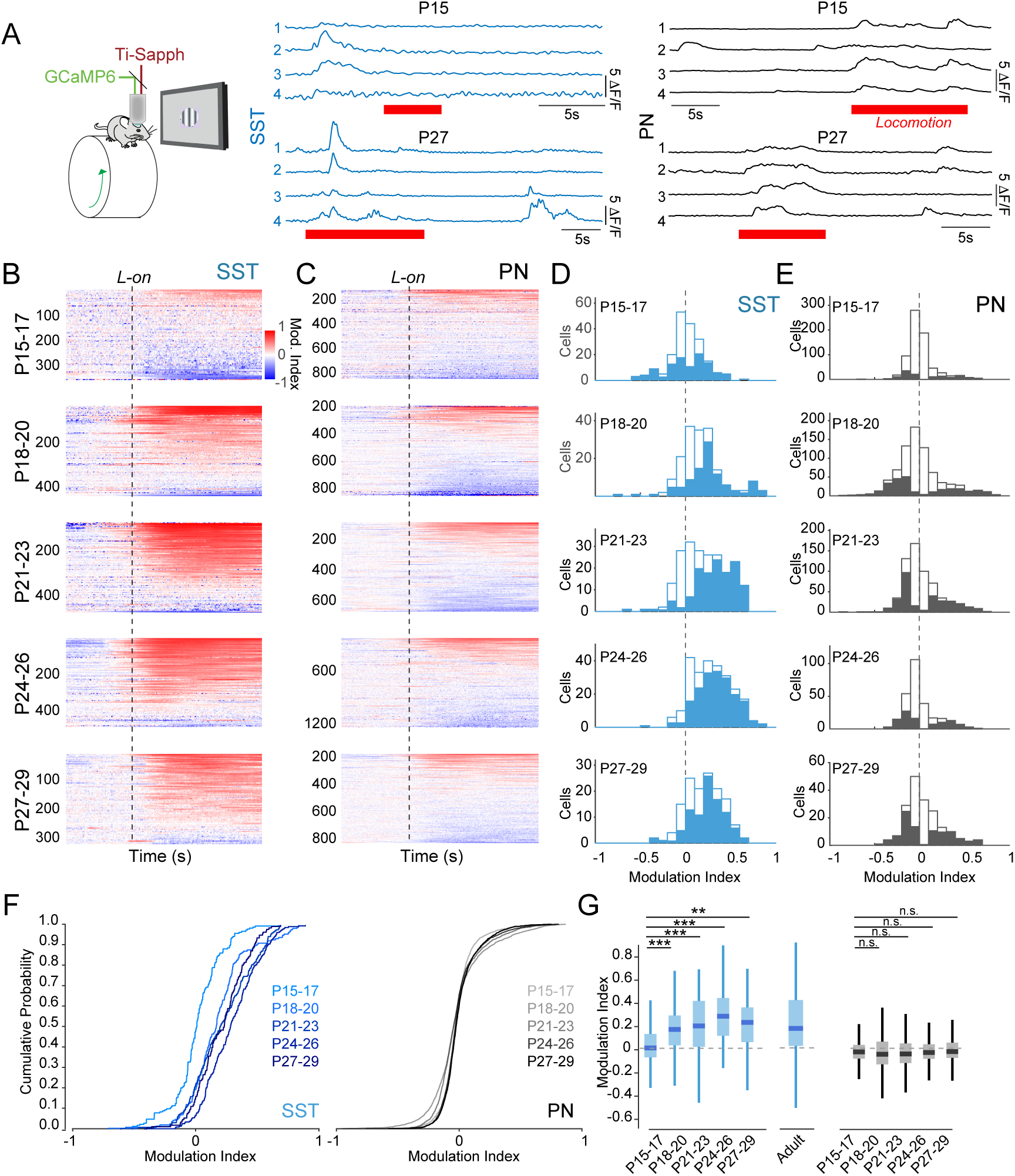
State-dependent modulation of SST-IN and PN activity across postnatal development. (A) Left: schematic of the in vivo 2-photon imaging configuration. Center: Ca2+ traces of four example SST-INs (blue) at P15 (upper) and P27 (lower). Right: Ca2+ traces of four example PNs (black) at P15 (upper) and P27 (lower). Locomotion bouts are indicated by red bars. (B) Modulation of activity around locomotion onset (L-on), calculated as an index value, for SST-INs in each age group. Each line represents the activity of a single cell exhibiting positive (red) or negative (blue) modulation. (C) Same as in (B) but for PNs. (D) Histograms of modulation indices of all SST-INs in each age group (P15-17: n = 197 cells, 6 mice; P18-20: n = 180 cells, 8 mice; P21-23: n = 211 cells, 9 mice; P24-26: n = 246 cells, 10 mice; P27-29: n = 142 cells, 6 mice). Solid bars indicate cells showing significant modulation at p <0.05 (shuffle test). (E) Same as in (D) but for PNs (P15-17: n = 3301 cells, 4 mice; P18-20: n = 2545 cells, 4 mice; P21-23: n = 2190 cells, 6 mice; P24-26: n = 3638 cells, 8 mice; P27-29: n = 2267 cells, 3 mice). (F) Cumulative probability distribution of locomotion modulation index for each age group of SST-INs (left) and PNs (right). (G) Boxplots of locomotion modulation indices across ages from (D) and (E) for SST-INs (blue) and PNs (black). SST-IN boxplots include an adult (>P150) index value for comparison. Central mark indicates the median and whiskers indicate 25th and 75th percentiles. *p<0.05, **p<0.01, ***p<0.001, linear mixed-effects model with age as fixed effect and mouse as random effect. Adult data are replotted from Ferguson et al. (2023).

**Supplemental Figure 3.**
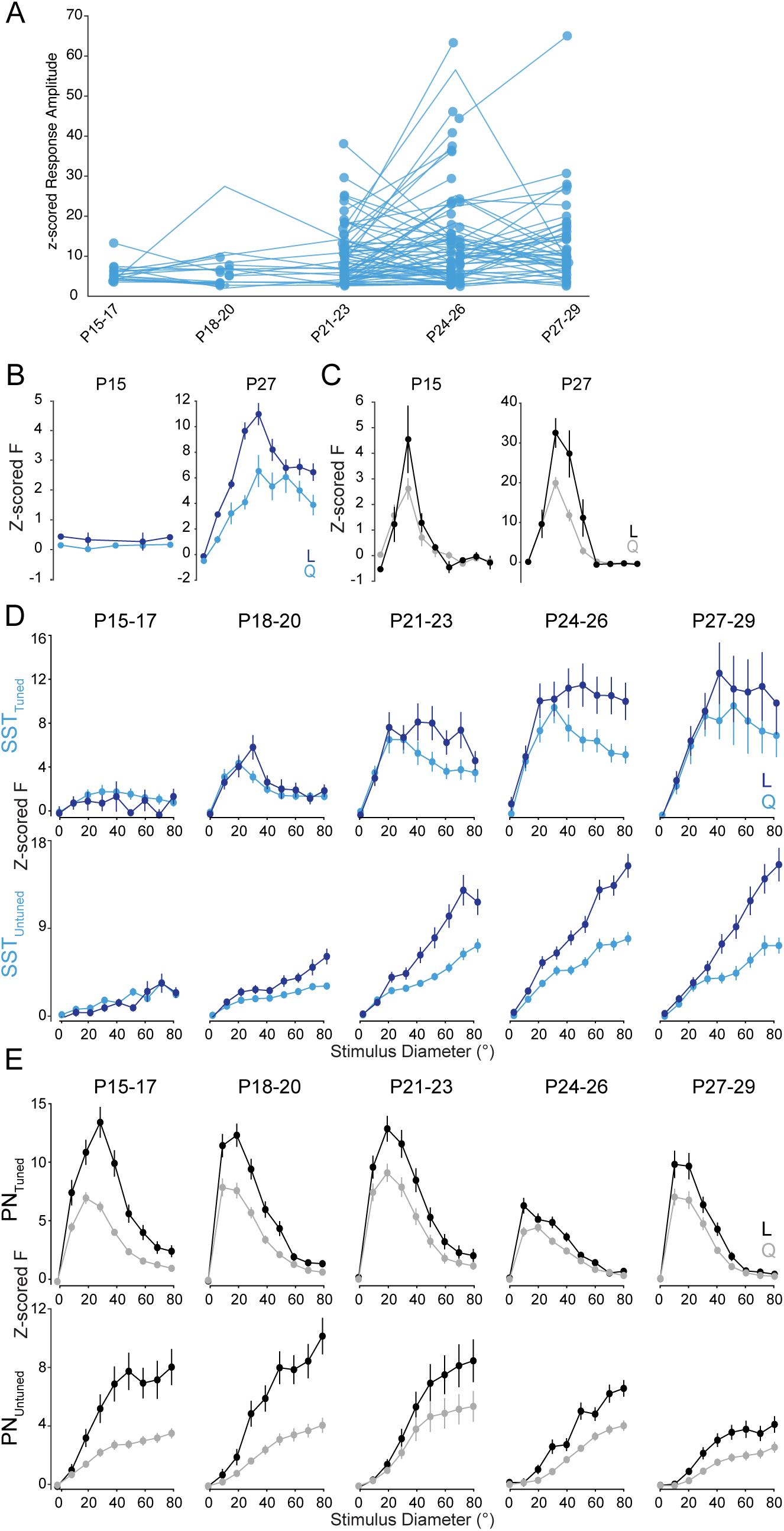
SST-IN visual response amplitudes increase after P15 and stabilize before P30. (A) (A) Visual response amplitudes of individual SST-INs imaged across days spanning age groups. Connected lines indicate response amplitudes measured from the same cell at different time points. (B) Mean visual responses of example P15 (left) and P27 (right) SST-INs to drifting grating stimuli of varying size. Responses are Z-scored to the 1-second baseline period before the stimulus onset for periods of quiescence (Q, light lines) and locomotion (L, dark lines). (C) Same as in (B) but for example PNs. (D) Visual response tuning curves for visually responsive SST-INs that were visually tuned (upper) or not tuned (lower) for stimulus size (see Methods) at each age during periods of quiescence (Q, light lines) and locomotion (L, dark lines). (E) Same as in D but for PNs.

**Supplemental Figure 4.**
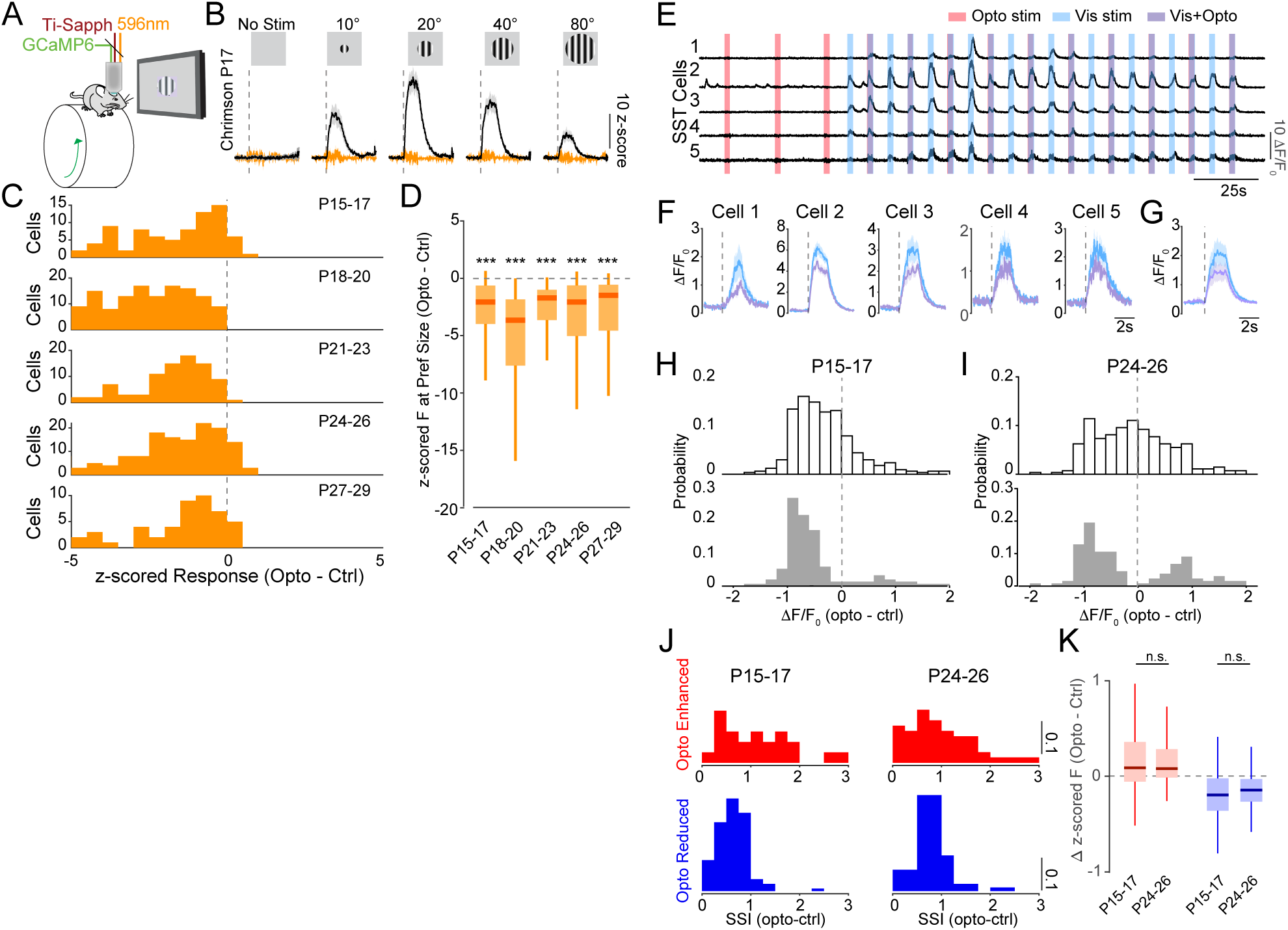
Inhibition of PNs by SST-INs throughout the P15-P29 period. (A) Schematic of experimental configuration for simultaneous in vivo optogenetics and 2-photon imaging. (B) Example P17 PN in an SST^Cre+^;Thy1-GCaMP6 animal expressing Cre-dependent Chrimson in SST-INs and GCaMP6 in PNs, showing visual responses to drifting grating stimuli of varying sizes during baseline conditions (black) and optogenetic activation of SST-INs (orange). Vertical dashed lines indicate visual stimulus onset. Shaded areas indicate mean ± SEM. (C) Histograms of the difference in z-scored response at preferred stimulus size between control and optogenetic trials for Chrimson-expressing mice across age groups (P15-17: n = 124 cells, 6 mice; P18-20: n = 268 cells, 7 mice; P21-23: n = 133 cells, 3 mice; P24-26: n = 264 cells, 6 mice; P27-29: n = 75 cells, 1 mouse). (D) Boxplots of the values in (C). Central mark indicates the median and whiskers indicate 25th and 75th percentiles. P15-17: n = 124 cells, 6 mice; P18-20: n = 268 cells, 7 mice; P21-23: n = 133 cells, 3 mice; P24-26: n = 264 cells, 6 mice; P27-29: n = 75 cells, 1 mouse. *p<0.05, **p<0.01, ***p<0.001, intercept-only linear mixed effects regression model with no fixed effect and mouse as random effect. (E) Example Ca2+ traces from 5 SST-INs in an SST^Cre+^;Ai148^F/0^ mouse expressing Cre-dependent ArchT. Red bars denote optogenetic stimulation, blue bars denote visual stimulation, and purple bars denote coincident optogenetic and visual stimuli. (F) Averaged visual responses from the cells shown in panel E. Responses to visual stimuli alone are shown in blue and responses to visual stimuli during optogenetic suppression with ArchT are shown in purple. (G) Population average of visual responses without (blue) and with (purple) optogenetic suppression via ArchT in all cells (n = 18) from the experiment shown in panels E and F. (H) Upper: Histogram of the difference in z-scored response at the preferred stimulus size between control and optogenetic trials for all PNs recorded in ArchT-expressing SST^Cre+^;Thy1-GCaMP6 animals at P15-17. Lower: Histogram of the subset of P15-17 PNs whose responses were significantly modulated. (I) Same as H, for P24-26. (J) Histograms of the surround suppression index values for the change in response amplitude in population of PNs exhibiting significantly enhanced (red) and reduced (blue) visual responses during SST-IN suppression at ages P15-17 (left) and P24-26 (right). (K) Impact of optogenetic suppression of SST-INs on spontaneous activity of PNs at P15-17 and P24-26 in PNs showing enhanced (red) and reduced (blue) responses. Enhanced PNs: P15-17: n = 202 cells, 6 mice; P24-26: n = 193 cells, 4 mice. Reduced PNs: P15-17: n = 166 cells, 6 mice; P24-26: n = 124 cells, 4 mice. Linear mixed-effects regression model with age as fixed effect and mouse as random effect.

